# Limited Volumetric Separation Across CDR Groups in OASIS-1

**DOI:** 10.64898/2026.05.27.728320

**Authors:** Zimmerman Kendra, Mahajan Shaurya, Sayadyan Diana, Peralta Roman, Tameze Paulyn, Gonzalez Matthew, Oushana Luka, Thunga Sanjay, Noah St. Clair

## Abstract

Clinical Dementia Rating (CDR) scores are used to classify the cognitive state of patients and are provided within neuroimaging datasets. This is achieved through a standardized clinical assessment that evaluates participants’ cognitive and functional abilities in everyday life, after which they are given a score ranging from 0 to 3. Where 0 represents no signs of dementia and three represents severe dementia^1^. These scores are then used to track the progression of dementia over time^2^. This study explored if these CDR labels within the OASIS-1 dataset produced consistent volumetric separation across the hippocampus, amygdala, and cortex.

## Introduction

This study examined whether CDR labels in the OASIS-1 MRI dataset correspond to consistent, significant patterns of brain atrophy. Many studies treating clinical labels, such as CDR 0, 0.5, and 1, are used to suggest cognitive decline relating to dementia. However, structural MRI differences between CDR groups were limited across several common neuroanatomical markers, including the hippocampus, amygdala, and cortex. This study analyzed structural MRI differences across CDR scores using hippocampus, amygdala, and the cortical volumetric measurements.

## Methods

Data was collected from the Open Access Series of Imaging Studies (OASIS-1) cross sectional MRI dataset. Participants included in this analysis were grouped according to their Clinical Dementia Rating (CDR) scores of 0, 0.5, or 1. Subjects with missing MRI volumetric measurements or incomplete CDR information were excluded from the analysis. The total cohort consisted of 73 individuals, 41 with CDR 0, 25 with CDR 0.5 and 7 with CDR 1. The hippocampus, amygdala, and cortex were selected as phenotypic markers of dementia. Left and right hemisphere measurements for the hippocampus and amygdala were combined to generate bilateral regional volumes. Cortical measurements were obtained using the total cortical gray matter volume provided in the dataset. To account for inter-subject differences in head size, all regional volumes were normalized to estimated total intracranial volume (eTIV). Pairwise comparisons were performed for each group. Because of unequal group sizes and non-normal distribution patterns observed within several MRI variables, nonparametric Mann Whitney U tests were used to compare normalized regional volumes between groups. Effect sizes were calculated using Cohen’s d to estimate the magnitude of volumetric differences between CDR groups. Statistical significance was defined as a p value < 0.05.

## Results

**Figure 1.**
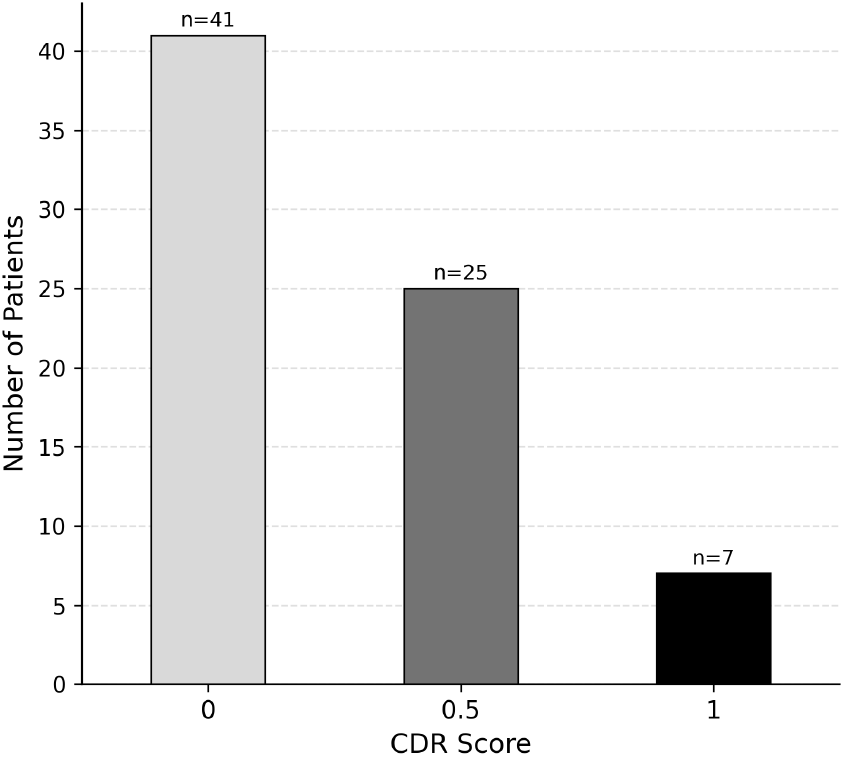
Distribution of patients with varying Clinical Dementia Ratings. Per the provided Oasis CDR scores, patients were grouped into CDR 0, 0.5, or 1. The sample contained 41 participants with CDR 0, 25 participants with CDR 0.5, and 7 participants with CDR 1.

**Figure 2.**
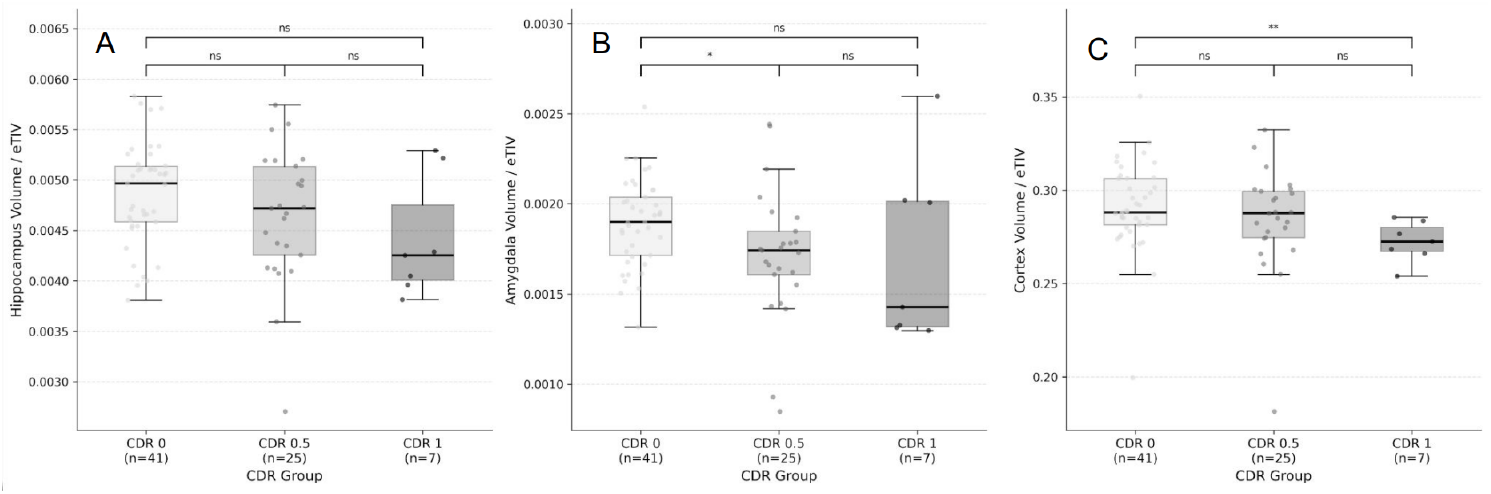
Boxplots showing the regional atrophy differences normalized to whole brain volume across CDR 0, 0.5, 1. Individual points represent different subjects. A) Hippocampal volume normalized to eTIV. B) Amygdala volume normalized to eTIV. C) Cortical volume normalized to eTIV. Pairwise statistical comparisons are shown above each plot, with “ns” indicating a nonsignificant comparison.

**Table 1.** Pairwise statistical comparison of normalized regional brain volumes between CDR groups.

| Comparison | Hippocampus<br>d | Hippocampus<br>p | Amygdala<br>d | Amygdala<br>p | Cortex<br>d | Cortex<br>p |
| --- | --- | --- | --- | --- | --- | --- |
| 0 → 0.5 | 0.39 | 0.245 | 0.59 | 0.021 | 0.25 | 0.348 |
| 0.5 → 1 | 0.35 | 0.281 | 0.02 | 0.624 | 0.47 | 0.054 |
| 0 → 1 | 0.88 | 0.091 | 0.62 | 0.225 | 0.84 | 0.003 |

Hippocampal volume was reduced in patients with higher CDR scores, with the CDR 1 group showing a smaller volume than the CDR 0 group. Amygdala volume showed a similar pattern: the CDR 0 group had a slightly larger volume than the CDR 0.5 group, while the CDR 1 group showed the greatest reduction overall. Cortex volume remained relatively consistent between the CDR 0 and CDR 0.5 groups; however, patients classified as CDR 1 exhibited a substantially smaller cortical volume compared to the other groups.

When comparing patients with CDR scores of 0 and 0.5, the amygdala demonstrated the strongest statistically significant difference, with a p value of 0.021 and a moderate effect size (Cohen’s d = 0.59). In contrast, the hippocampus (p = 0.245, d = 0.39) and cortex (p = 0.348, d = 0.25) showed weaker differences between the two groups. In the comparison between CDR 0.5 and CDR 1, none of the analyzed brain regions reached statistical significance; however, the cortex produced the largest effect size (d = 0.47) along with a nearly significant p value (p = 0.054), suggesting greater cortical volume changes with increasing dementia severity. Comparing CDR 0 and CDR 1, the cortex displayed the most significant overall difference, with a low p value (p = 0.003) and a large effect size (d = 0.84). The hippocampus and amygdala also demonstrated large effect sizes, (d = 0.88) and (d = 0.62) respectively, despite not reaching statistical significance, (p = 0.091) and (p = 0.225) respectively, indicating substantial structural differences between non-demented and more severely demented patients.

## Discussion

The OASIS-1 dataset suggests limited volumetric separation across CDR groups. The majority of the CDR score comparisons resulted in p-values above the threshold for statistical significance, with the exception of the amygdala 0 - 0.5 CDR comparison giving a p-value of 0.021 and the cortex 0 - 1 CDR comparison giving a p-value of 0.003.

Looking more closely at the 0 - 0.5 CDR sample, this cohort signifies the early transition from healthy individuals to early signs of cognitive decline. Across the hippocampus, amygdala, and cortex, the results remained inconsistent between CDR groups and did not show clear patterns of brain atrophy.

The hippocampus did not suggest significant differences between any of the CDR groups. The comparisons between CDR 0 and 0.5 produced a small effect size (d=.39, p=.245), while the comparison between CDR 0.5 and 1 also remained insignificant (d=.35, p=.281). Although the comparison between CDR 0 and 1 produced a large effect size (d=.88), the result still failed to reach statistical significance (p=.091). Hippocampal atrophy is a well-established marker of dementia, making the lack of significant differences across comparisons notable within the dataset^3^. The large effect size between CDR 0 and 1 without statistical significance may reflect limited statistical power. Additionally, the CDR 1 group contained a small sample size (n=7), potentially limiting the strength of the comparison.

For the amygdala, the comparison between CDR 0 and 0.5 shows a moderate effect size and is the only significant comparison in the entire dataset (d=.59, p=.021). However, the comparisons between CDR 0.5 and 1 shows almost no effect ( d = .02, p=.624). The comparison between CDR 0 and 1 shows a moderate effect size (d=.62, p=.225) but it is not significant. The sharp drop from d=.59 to d=.02 across CDR progression further supports variability across CDR comparisons.

For the cortex, the comparison between CDR 0 and 0.5 showed an insignificant effect (d=.25, p=.348), while the comparison between CDR 0.5 and 1 showed a moderate effect that was still not strongly significant (d=.47, p=.054 ). The comparison between CDR 0 and 1 produced a large effect size that was significant (d=.84, p=.003), making the cortex the only region with a clearly significant effect. However, the CDR 0.5 group was not significantly different from either CDR 0 (p=.348) or CDR 1 ( p=.054). Although the effect sizes trend in the expected direction from .25 to .47 to .84, the significance does not follow the same progression.

Several limitations should be considered when interpreting these findings. The CDR 1 cohort contained only seven participants, limiting statistical power and increasing variability across comparisons. Additionally, only a limited set of volumetric MRI measures were analyzed, and therefore these findings should not be interpreted as representing all neuroanatomical features within the dataset. Because analyses were exploratory, multiple comparison correction was not applied and isolated significant findings should be interpreted cautiously.

Overall, the analyzed regions showed inconsistent separation across CDR groups. There are a few isolated significant findings, specifically the amygdala (0 - 0.5) and cortex (0 - 1), but these findings were not consistently replicated across the remaining comparisons. Additionally, the CDR 1 group contains a small sample size (n=7), which limits its ability to be able to support any strong conclusions even at the most severe stage.

## Footnotes

1 Wang G, Li Y, McDade E, et al. Clinical progression on CDR-SB©: Progression-free time at each 0.5 unit level in dominantly inherited and sporadic Alzheimer’s disease populations. Alzheimer’s Dementia.

2 Marcus, D. S., Fotenos, A. F., Csernansky, J. G., Morris, J. C., & Buckner, R. L. (2010). Open Access Series of Imaging Studies (OASIS): Longitudinal MRI Data in Nondemented and Demented Older Adults. *Journal of Cognitive Neuroscience, 22*(12), 2677–2684.

3 Schuff, N., Woerner, N., Boreta, L., Kornfield, T., Shaw, L. M., Trojanowski, J. Q., Thompson, P. M., Jack, C. R., Jr, Weiner, M. W., & Alzheimer’s Disease Neuroimaging Initiative (2009). MRI of hippocampal volume loss in early Alzheimer’s disease in relation to ApoE genotype and biomarkers. *Brain* : *a journal of neurology, 132*(Pt 4), 1067–1077.

